# A role for root carbonic anhydrase βCA4 in bicarbonate tolerance of *Arabidopsis thaliana*

**DOI:** 10.1101/2024.04.18.590072

**Authors:** Laura Pérez-Martín, Maria-José Almira, Laura Estrela-Muriel, Roser Tolrà, Lourdes Rubio, Charlotte Poschenrieder, Silvia Busoms

**Author notes:** **Author for correspondence:** Silvia Busoms. Contributed equally (Laura Pérez-Martín, Maria-José Almira).

## Abstract

Carbonic anhydrases (CAs) are the main enzymes handling bicarbonate in the different cell compartments. This study analyses the expression of CAs in roots of *Arabidopsis thaliana* demes differing in tolerance to bicarbonate: the tolerant A1_(c+)_ deme and the sensitive deme, T6_(c-)_. While 10 mM NaCl caused a transient depolarization of the root cell membranes, 10 mM NaHCO_3_ caused hyperpolarization. This hyperpolarization was much stronger in A1_(c+)_ than in T6_(c-)_. Acetazolamide (AZ), a specific inhibitor of CAs, abolished the hyperpolarizing effect in A1_(c+)_, indicating the implication of CAs in this fast membrane response. The time dependent (3 to 72 h) expression profiles of 14 CAs (αCA1-8 and βCA1-6) in roots of A1(c+) and T6_(c-)_ exposed to either control or NaHCO_3_ (pH 8.3) revealed a bicarbonate specific upregulation of βCA4.1 (from 3 to 12 h) and, although to a lesser extent, of βCA3 in A1_(c+)_. Contrastingly, in T6_(c-)_ βCA4 was downregulated by NaHCO_3_. Exclusively in A1_(c+)_, the enhanced expression of βCA4 under bicarbonate was parallelled by an increase of *PIP1,3, SLAH1, SLAH3, AHA2*, and *FRO2* gene expression levels. Under HCO_3_ ^-^ exposure, a *βca4* knockout mutant had lower number of lateral roots, lower root diameter and higher MDA root concentrations than the WT. The obtained results indicate that bicarbonate induced root membrane hyperpolarization is the fast (minutes) initial signalling event in the tolerance response, followed by the specific upregulation of β*CA4*.*1* and the genes involved in H20 and CO_2_ transport, apoplast acidification, ion homeostasis and iron acquisition.

## 1 Introduction

Alkaline soils currently cover more than 30% of the earth surface, especially in arid and semiarid areas such as the Mediterranean (Wahba et al., 2019). Under the prevailing and future climate change scenarios an aggravation of crop land alkalinization is foreseen (Hassani et al., 2021) and breeding for alkaline tolerant crops is an urgent mission.

Limestone soils, alkaline and alkaline saline soils largely differ in their physical and chemical properties and the diverse constraints they impose on crop performance are increasing as pH and sodicity build up. However, a common characteristic is the high concentration of bicarbonate in their soil solutions.

The adverse effects of bicarbonate on sensitive crop plants have mainly been considered by its repercussion on iron deficiency and, to a lesser extent on other mineral nutrient deficiencies like nitrogen (N), potassium (K), magnesium (Mg), Zinc (Zn), manganese (Mn), molybdenum (Mo), copper (Cu) and zinc (Zn) deficiency. Chalk chlorosis is a typical symptom of bicarbonate-induced iron deficiency in calcifuge species cultivated on calcareous soils (Tagliavini and Rombolà, 2001). Several mechanisms are responsible for this Fe deficiency: low Fe availability in soils with alkaline pH, interference of carbonate/bicarbonate with the iron mobilization strategy 1 of dicot species, hampering the activity of FRO2, the iron reductase that requires acid pH for optimal function, or the inhibition of Fe transport from roots to shoots (Wang et al., 2020).

While carbonate/bicarbonate by its buffer capacity may efficiently counteract the Fe acquisition strategy based on rhizosphere acidification, the exudation of flavonoid–type phenolics able to chelate Fe even under alkaline pH conditions is a major adaptive mechanism for plant performance on carbonate–rich soil (Terés et al., 2019; Vélez-Bermudez and Schmidt, 2023). Fe deficiency responses on the genetic and molecular level has attracted much research effort and the basic mechanisms of Fe acquisition and transport are getting highly established (Riaz and Guerinot, 2021), however the fate of bicarbonate in plants is more concealed (Poschenrieder et al., 2018).

Carbonic anhydrases (CAs) are the main enzymes handling bicarbonate both inside and outside plant cells (DiMario et al., 2017, 2018; Ignatova et al., 2019). The role of CA in carbon concentration mechanisms in aquatic photosynthetic organisms is well established (Raven et al., 2011; Kupriyanova et al., 2023). Leaf CAs have recently gained much attention not only in C4 photosynthesis (Zhou et al., 2024) but also in terrestrial C3 plants due to their potential role in mesophyll conductance and photosynthesis, especially under stress conditions (Momayezzi et al., 2019, Weerasooriya et al., 2024). In carbonate-rich karst environments drought-induced stomatal closure may strongly limit CO_2_ diffusion into the leaves. In plants adapted to such extreme conditions, the overall photosynthesis rate is considerably decreased, but CA activity is strongly enhanced and HCO_3_ ^-^ -derived CO_2_ can contribute up to 30% of the photosynthetic C assimilation (Wu and Xing, 2012; Wu and Wu, 2022). The ability to enhance CA in response to HCO_3_ ^-^ exposure has been highlighted as an important trait for adaptation to the karst environment (Müller et al., 2014)

Nonetheless, the information on the mechanisms behind the potential role of root CAs in the tolerance of terrestrial plants to high soil bicarbonate is poor. The elucidation is hampered by the large number of different CAs located in different cell compartments in plants and the still fragmentary identification of the corresponding genes in bicarbonate tolerant species. Best characterized from the genetic point of view is the model plant *Arabidopsis thaliana*. The species contains 19 CA genes: 8 α, 6 β, and 5 gamma CAs (DiMario et al., 2018).

*Arabidopsis thaliana* is increasingly employed in evolutionary and ecological studies. Patterns of natural genetic variation and the forces that shape them have been studied mostly at global and regional scales (Stenøien et al., 2005; Schmid et al., 2006; Beck et al., 2008). Local populations have shown to be strongly differentiated even when they are geographically close (Bergelson et al., 1998; Bakker et al., 2006), proving the effects of environmental heterogeneity in creating diversity patterns even at local scale. Genotypic and phenotypic diversity of natural subpopulations (demes) of *A. thaliana* from the North East of Spain (Catalunya) were large enough to provide an scenario for exploring tolerance mechanisms to soil carbonate (Terés et al., 2017, Pérez-Martín et al., 2021): two demes, A1_(c+)_ and T6_(C-)_ with differential tolerance to moderate soil carbonate concentrations showed differential growth and reproduction and photosynthetic efficiency due to differences in both nutrient profiles and degree of membrane damage under bicarbonate treatment (Pérez-Martín et al., 2021).

With the aim of shedding light on the role of CAs in plant bicarbonate tolerance and taking advantage of these naturally evolved differences in tolerance to carbonated soils, the present study explores the influence of bicarbonate on the plasma membrane potential of root cells and the expression levels of α and β CAs in roots of these contrasting *A. thaliana* demes. Our working hypothesis was that the tolerant A1_(C+)_ may better control bicarbonate toxicity by higher expression levels of carbonic anhydrases.

## 2 Materials and Methods

### 2.1 Plant materials

Seeds of contrasting phenotype lines from A1_(C+)_, an alkaline-moderate tolerant deme, and T6_(c-)_, an alkaline sensitive deme, were collected from field cultivated plants and stored under cold (4 ºC) and dry conditions until the beginning of the experiments. Col-0 seeds from Nottingham Arabidopsis Stock Centre (NASC) (Scholl et al., 2000) were included as a reference. Based on gene expression results, a mutant knock-out line (ecotype Columbia) carrying T-DNA insertions in βCA4, β*ca*4-1, (WiscDsLox508D11) obtained from Nottingham Arabidopsis Stock Centre (NASC) was included in hydroponic studies and the phenotypic analysis.

### 2.2 Growth Conditions

Seeds were surface sterilized by soaking in 30% Clorox bleach and a drop of Tween-20 for 10 minutes, and then rinsed 6 times in sterile 18 MΩ milli-Q water. Seeds were stored in KNO_3_ solution at 4 °C and darkness for stratification for 1 week.

For electrophysiological studies, seedlings were cultivated on plates. Seeds from each deme were sown in plates under a flow cabinet with sterile material. Plates contained control (½ MS at pH 5.9), Phyto-agar 0.6% (Duchefa, Haarlem, The Netherlands). Plates with seeds were kept at 4 °C for synchronizing germination. After 7 days under stratification treatment, plates were moved to a growth chamber (12 h light/12 h dark, 150 μmol cm^−2^•s^−1^, 40% humidity and 25 °C) where seedlings were grown for 7 further days.

For all other experiments *A. thaliana* plants were grown in hydroponics. For this purpose, seeds were sown in 0.2 mL tubes containing 0.6% agar prepared in nutrient solution 1/2 Hoagland (pH 5.9). Seeds were kept at 4ºC for 7 days in the dark to synchronize germination. Tubes containing seeds were then placed in the growth chamber under conditions described above. After root emergence, the bottoms of the tubes containing seedlings were cut off and the tubes were placed in 150 mL hydroponic containers with aerated ½ Hoagland solution (pH 5.9). 15 days old plants were separated in two different treatments: control (½ Hoagland solution at pH 5.9) and bicarbonate (½ Hoagland solution with 10 mM NaHCO_3_ at pH 8,3). Solutions were buffered with different proportions of MES and BTP depending on the desired final pH. The nutrient solutions were changed every 3 days.

### 2.3 Phenotypic analysis

After 10 days of treatment, the plants were harvested for phenotypic analysis. Mini-Pam II was used for measuring photosynthetic efficiency in leaves and leaf chlorophyll concentration was measured using Opti-Science CCM-300. Morphology was assessed by scanning the roots with an Epson Expression 10000XL scanner and analyzed using the software WinRhizo Pro 2009c. Average root length, number of lateral roots, number of forks and tips and average root diameter were analyzed.

For malondialdehyde (MDA) analysis, 200 μL of 0.5% thiobarbituric acid (TBA) in 20% trichloroacetic acid (TCA) was added to 200 μL of the extract. The mixtures were heated at 95 °C for 25 min and then quickly cooled in an ice bath. After centrifugation at 800 rpm for 6 min, the absorbance was recorded at 532 and 600 nm. Lipid peroxidation is expressed as nmol MDA x g^-1^ FW.

### 2.4 Root cell membrane potential

Plate cultivated, 7-day old seedlings were exposed to different NaCl or NaHCO_3_ concentrations ranging from 0 to10 mM in buffered solutions (pH 8.3). The influence of the inhibition of external carbonic anhydrases was studied using 10 mM acetazolamide (AZ) solution diluted in 0.05N NaOH (Rubio et al., 2017). The microelectrodes were produced using 1.5 mm OD glass capillaries pulled in a David Kopf vertical puller (model 720, David Kpoft Instruments, Tujunga, CA). The reference electrode was filled with 0.5 M KCl + agar 0.03% while the measurement electrode was filled with 0.5M KCl. Both electrodes were connected to a high-impedance differential amplifier (FD223a). The measurement chamber was open on both sides allowing the approach of several electrodes to the root. A gravity-based flow-through system permitted controlled changes of the medium at a rate of 10 ml min^-1^. This system kept the temperature, ionic concentration, and gases constant during the experiments. Microscope light (150 μmol photons m^-2^ s^-1^) was on during the experiments. The signals from differences in root membrane potential were monitored on a chart recorder (Linseis L250E).

### 2.5 Relative gene expression

Hydroponically grown 7 days old seedlings were transferred to treatment solutions. Root material was extracted after 0, 3-, 6-, 24-, and 48-hours treatment. Three plants were pooled to perform a replicate and 3 replicates were taken from each deme and treatment. Root material was rinsed with deionized water and immediately frozen at -80 ⍰C.

RNA was extracted using Maxwell plant RNA kit (Promega Corporation, Madison, WI, USA). After quality and quantity control, RNA was transformed to cDNA using iScriptTM cDNA Synthesis Kit (Bio-Rad, USA). Dilution of the cDNAs was performed 1/50 with water (Molecular Biology Reagent, Sigma-Aldrich, St. Louis, MO, USA). Diluted cDNA (1:50) was used as a template for quantitative PCRs using iTaq Universal SYBR Green Supermix (Bio-Rad, Hercules, CA, USA). Real-time detection of fluorescence emission was performed on a CFX384 Real-Time System (Bio-Rad, Hercules, CA, USA) using the following conditions: denaturalization step 10’’ 9511C followed by annealing and extension 30’’ 60⍰ C. A total of 40 cycles were run. A melting curve was performed, increasing from 65.0⍰ C to 95.0⍰ C by 0.5⍰ C each 5 seconds. Primers for FRO2, IRT1, AHA2, αCAs (1-8) and βCAs (1-6) were designed using NCBI primer blast tool and they were purchased from Biolegio (Nijmegen, The Netherlands). Actin was used as a housekeeping gene. Sequence of the primers are shown in Dataset S1. Plates were edited using the CFX manager version 3.1 software. The expression of the target gene relative to the expression of the reference gene was calculated using the 2 ^−ΔΔCt^ method (Livak & Schmittgen, 2001).

### 2.6 Statistical analysis

Statistics and data visualization performed using JMP Pro 13.2.1 (SAS Institute Inc., Cary, NC, USA). Significance of differences was determined using the Dunnett test. In physiological measurements, the normal distribution of data was confirmed by Levene’s test. Different treatments and populations were analysed by two-way ANOVA. Post-hoc analyses were realized using the Tukey test. The number of plants used in each experiment and the statistical analyses are provided in Dataset S2.

## 3 Results and Discussion

Both *A. thaliana* demes clearly differed in their responses to bicarbonate stress. PSII efficiency measured as Fv/Fm ratio was considerably more affected by bicarbonate (10 mM NaHCO_3_, pH 8,3) in deme T6_(C-)_ than in A1_(C+)_ or in the reference accession Col-0 (Figure 1A). Root MDA concentrations as assessed by the TBA/TCA assay revealed higher bicarbonate–induced lipid peroxidation in T6_(C-)_than in A1_(C+)_ (Figure 1B). These results confirm the previously reported bicarbonate sensitivity in T6_(C-)_ and the contrasting tolerance in A1_(C+)_ (Terés et al., 2019; Perez Martin et al 2021).

**Figure 1.**
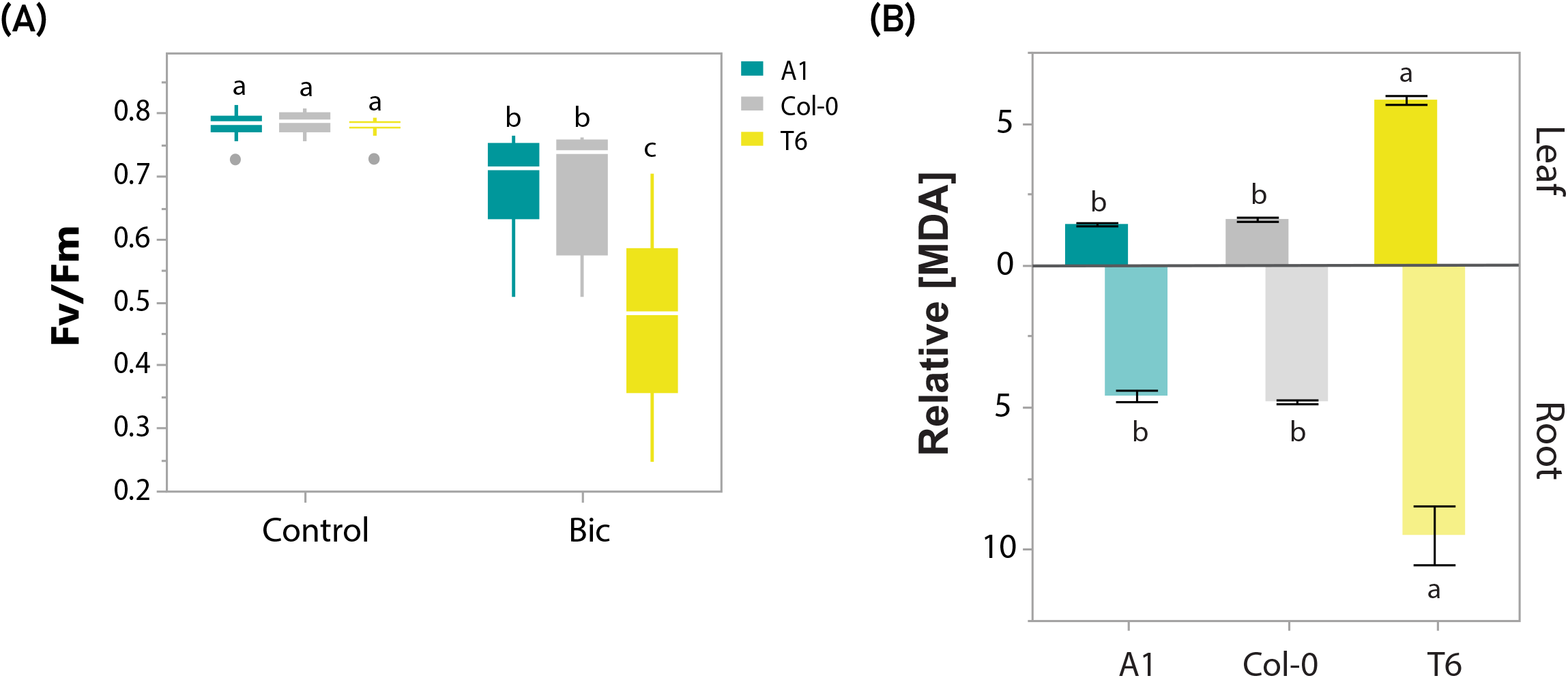
Photosynthesis efficiency and lipid peroxidation in Col-0, A1_(c+)_ and T6_(c−)_ under bicarbonate treatment. **(A)** Fv/Fm ratio (mean ± SE) of the reference accession Col-0, the tolerant A1_(c+)_ and the sensitive T6_(c−)_ demes under Control and Bic conditions (N=6). **(B)** Leaf (above) and root (below) relative MDA content [(MDA_Bic_/Mean MDA_C_); mean ± SE] of Col-0, A1_(c+)_ and T6_(c−)_ (N=3). Mean values with different letters indicate significant differences (Tukey’s HSD, adj. p-value < 0.05).

### 3.1 Root cells plasma membrane potential

To further characterizing the ion specificity of the observed differences, we compared the influence of NaCl and NaHCO_3_ on the root cell plasma membrane potential of both demes at isomolar concentrations and under the same alkaline pH (8.3) conditions (Figure 2A). Figures 2 a and b show the mean values of plasma membrane potential changes in epidermal root cells of A1_(C+)_ and T6_(C-)_ exposed to different NaCl and NaHCO_3_ concentrations, respectively. As expected, NaCl caused membrane depolarizations, that were higher in the case of T6 at higher concentration. The NaCl-induced depolarizations in A1_(C+)_ adjusted to a saturation curve showing a semisaturation constant (3.1± 0.7 mM NaCl) in the range of assayed NaCl concentrations while this was not the case in T6_(C-)_ (Figure 2A). Contrastingly, equimolar NaHCO_3_ exposure quickly caused a strong, transient hyperpolarization in A1_(C+)_, while less hyperpolarization was observed in T6_(C-)_ (Figure 2B). The different response caused by NaCl and NaHCO_3_ clearly demonstrate an anion specific effect. It is well established that under NaCl stress, Na^+^ uptake into root cells causes a quick transient depolarization of the membrane. In consequence, K^+^ efflux channels like GORK are activated leading to K^+^efflux, membrane repolarization and loss of cytosolic K^+^ homeostasis in sensitive plants, while salt tolerant plants are able to maintain K^+^ homeostasis (Wu et al., 2018).

**Figure 2.**
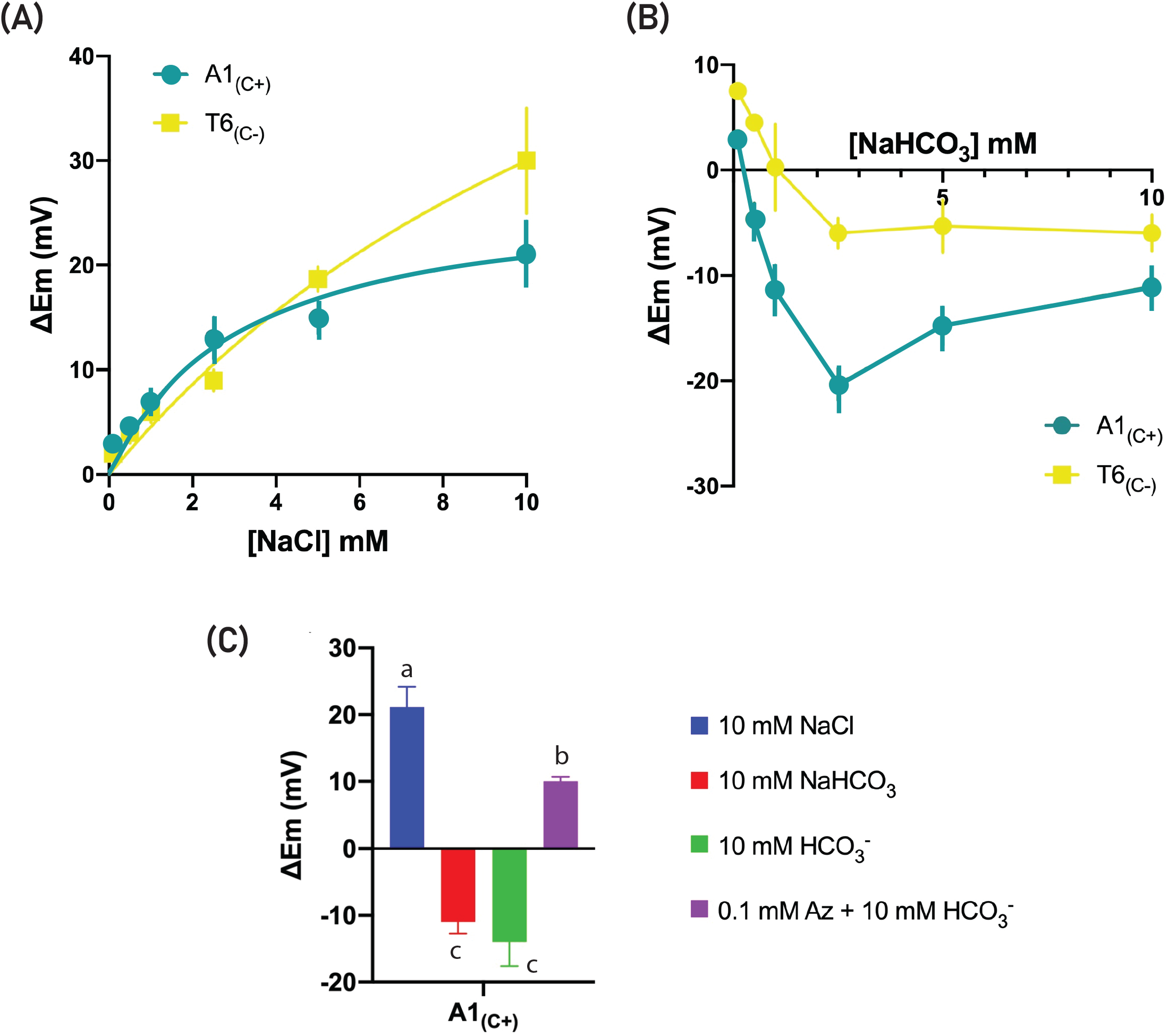
Effect of NaCl or NaHCO_3_ additions on plasma membrane potential of epidermal root cells of A1_(c+)_ and T6_(c−)_. **(A)** Plasma membrane depolarizations induced by increasing NaCl concentrations. Values showed saturation kinetics and were fitted to Michaelis-Menten model. **(B)**. Plasma membrane hyperpolarizations induced by increasing NaHCO_3_ concentrations. **(C)**. Comparison of plasma membrane variations in response to the addition of 10 mM NaCl, 10 mM NaHCO_3_, and 10 mM HCO_3_ ^-^ in the absence or in the presence of 0.1 mM acetazolamide (AZ). Data are mean± of at least five independent measurements. Different letters indicate significant differences (One way ANOVA, Tukey’s test, adj. p-value <0.0001).

Contrastingly, when Na^+^ was given in the form of NaHCO_3_, membrane hyperpolarization occurred (Figure 2B). NaHCO_3_ -induced hyperpolarization were higher in A1 than in T6 at external concentration from 1 to 10 mM. Hyperpolarization can be caused either by cation efflux or anion influx. To test whether bicarbonate uptake may be responsible for the strong hyperpolarization in A1, the influence of AZ, an inhibitor of carbonic anhydrases, on the bicarbonate-induced membrane hyperpolarization was tested. Since the addition of NaHCO_3_ would simultaneously promote a (Na^+^ -induced) membrane depolarization and a membrane hyperpolarization, if HCO _-_ is incorporated, we analyse the effect of the change of 10 mM NaCl by 10 mM NaHCO_3_ during the treatment in the absence or in the presence of 0.1 mM AZ. The presence of AZ completely abolished the hyperpolarizing effect of HCO_3_ ^-^ (Figure 2C). Carbonic anhydrases catalyse the reversible hydration of CO_2_ to HCO_3_ ^-^ and H^+^. Under the alkaline pH conditions of our experimental solution (pH 8.3) the supplied NaHCO_3_ was mainly in the form of HCO_3_ ^-^ and the CO_2_ concentration was negligible. In the more acidic pH values of the apoplast and the cytosol the proportion of CO_2_ should be considerably higher. At present, no specific bicarbonate transporters have been identified in higher terrestrial plants, but HCO_3_ ^-^ may enter cells by anion transporters (Poschenrieder et al., 2018). Bicarbonate entering the symplasm would cause strong hyperpolarization. However, the cancellation by the inhibition of CAs indicates that the observed membrane hyperpolarization was most probably caused by bicarbonate generated by the activity of carbonic anhydrase inside the cells. Another possibility is that the activity of carbonic anhydrases activates proton extrusions, leading to membrane hyperpolarization. The consequent acidification of the apoplast would favour HCO_3_ ^-^ dehydration and diffusion of CO_2_ into the cell, where internal CAs may convert it to HCO_3_ ^-^ contributing to further membrane hyperpolarization. Future experiments analysing cytosolic and external pH changes associated with this membrane hyperpolarization are required to clarify the underlying processes.

Regardless the mechanism, an interesting result is that the CA-induced membrane hyperpolarization was much stronger in A1_(C+)_ than in T6_(C-)_ (Figure 2B). Such a fast differential response in the tolerant A1_(C+)_ suggests that this membrane hyperpolarization could be the trigger of a signal transduction pathway leading to the quick activation of tolerance mechanisms in the A1_(C+)_ deme. In fact, strong root cell hyperpolarization can activate Ca^2+^ influx channels in *A. thaliana* (Kiegle et al., 2000; Demichick et al., 2002) thus initiating stress signalling and activation of tolerance related genes.

### 3.2 Relative expressions of root α- and β CAs and Fe-deficiency related genes

To explore this possibility, and following our initial hypothesis of a role for CAs in bicarbonate tolerance, we analysed the time-dependent root expressions of genes coding for α- and β CAs, the main responsible for bicarbonate handling in the apoplast, plasma membrane and cytosol (DiMario et al., 2017) and AHA2, FRO2, and IRT1 as the key players in root iron acquisition (Martín-Barranco et al., 2020). Figure 3 shows the heat map of root α- and β CA relative expressions in A1_(C+)_ and T6_(C-)_ between 3 to 72 h after bicarbonate exposure. The most conspicuous differences between both demes were the strong upregulation of βCA4 during the first hours in the tolerant A1_(C+)_, while the expression was downregulated in the sensitive T6_(C-)_.

**Figure 3.**
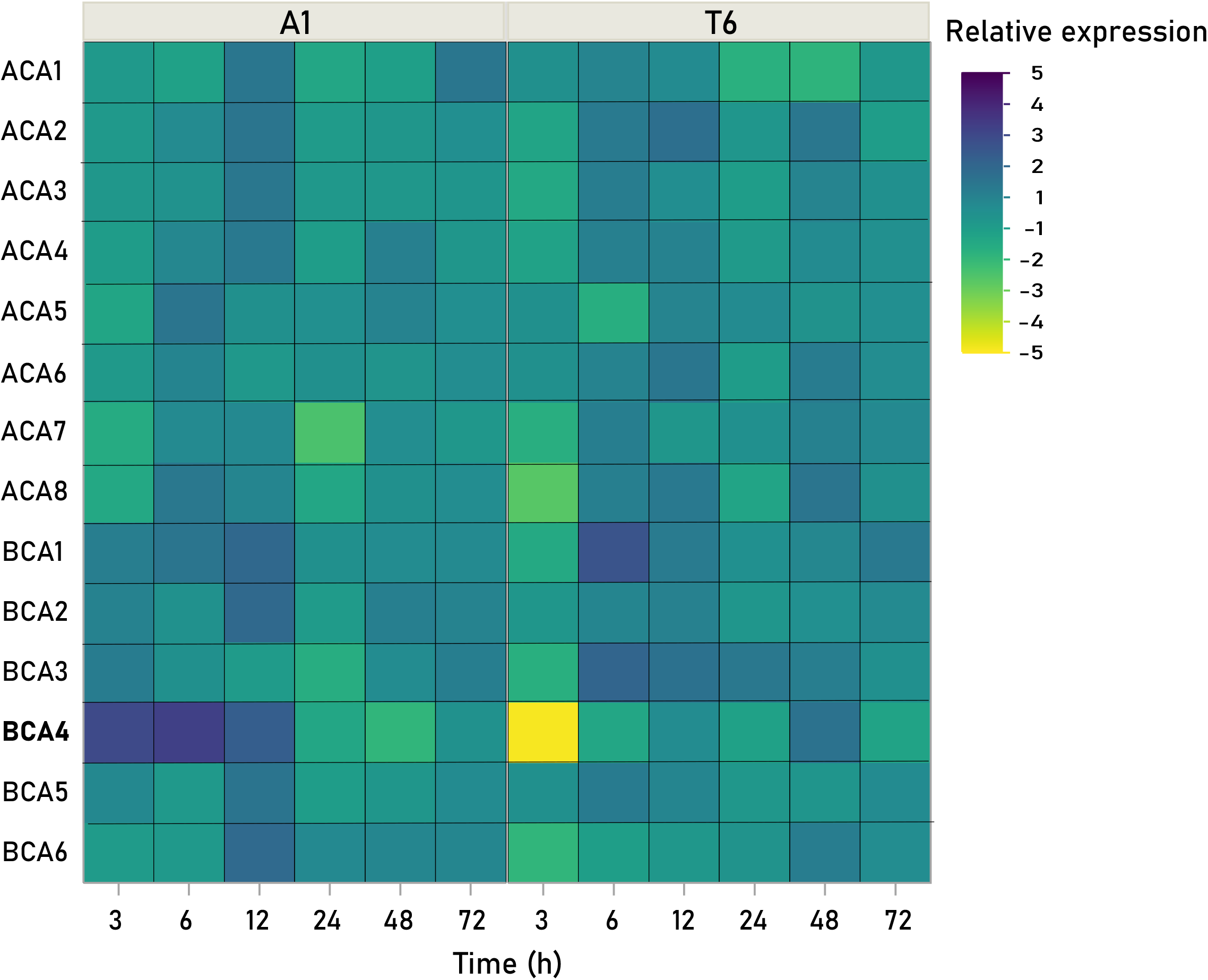
Root expression profiles of α- and β CAs. Gene-expression heatmap of α- and β Carbonic Anhydrases relative expression (log2(FC)_Bic_ / log2(FC)_C_) in roots of A1_(C+)_ and T6_(C-)_ after 3 or 72 h of bicarbonate exposure. The colour scale indicates the expression levels (dark blue, high relative expression; yellow, low relative expression).

In *A. thaliana* βCA4 is encoded by At1G70410 (βCA4) which presents 3 splice variants. The primer used in our experiment amplifies At1G70410.2 coding for βCA4.1 which is localized in the plasma membrane (DiMario et al., 2016). The role of plasma membrane bound CAs in roots has not been identified so far. In stomata the plasma membrane associated βCA4 mediates the control of stomatal movement by CO_2_ (Hu et al, 2015). βCA4–induced increase of HCO_3_ ^-^ concentration in the guard cells activates S-type anion channel SLAC1 promoting anion efflux and stomatal closure (Tian et al., 2015).

Relative expression profiles of selected genes related to iron acquisition are shown in Figure 4. In A1_(C+)_ the expressions of both *AHA2* (Figure 4A) and *FRO2* (Figure 4B) were upregulated by bicarbonate between 3 to 24 and 3 to 48 hours, respectively and declined thereafter. In T6_(C-)_ *AHA2* expression remained unchanged upon bicarbonate exposure and *FRO2* expression was initially decreased and increased only slightly after 12 to 48 h exposure. *IRT1* expression peaked earlier in T6_(C-)_ (6 to 12 h) than in A1_(C+)_ (24 to 48 h) (Figure 4C). AHA2 is the key player in apoplastic and rhizosphere acidification required for Fe mobilization and the activity of FRO2, the ferric chelate reductase. The quick upregulation of *AHA2* and *FRO2* prior to I*RT1* upregulation clearly support the superior ability of A1_(C+)_ to mobilize and reduce Fe under alkaline conditions.

**Figure 4.**
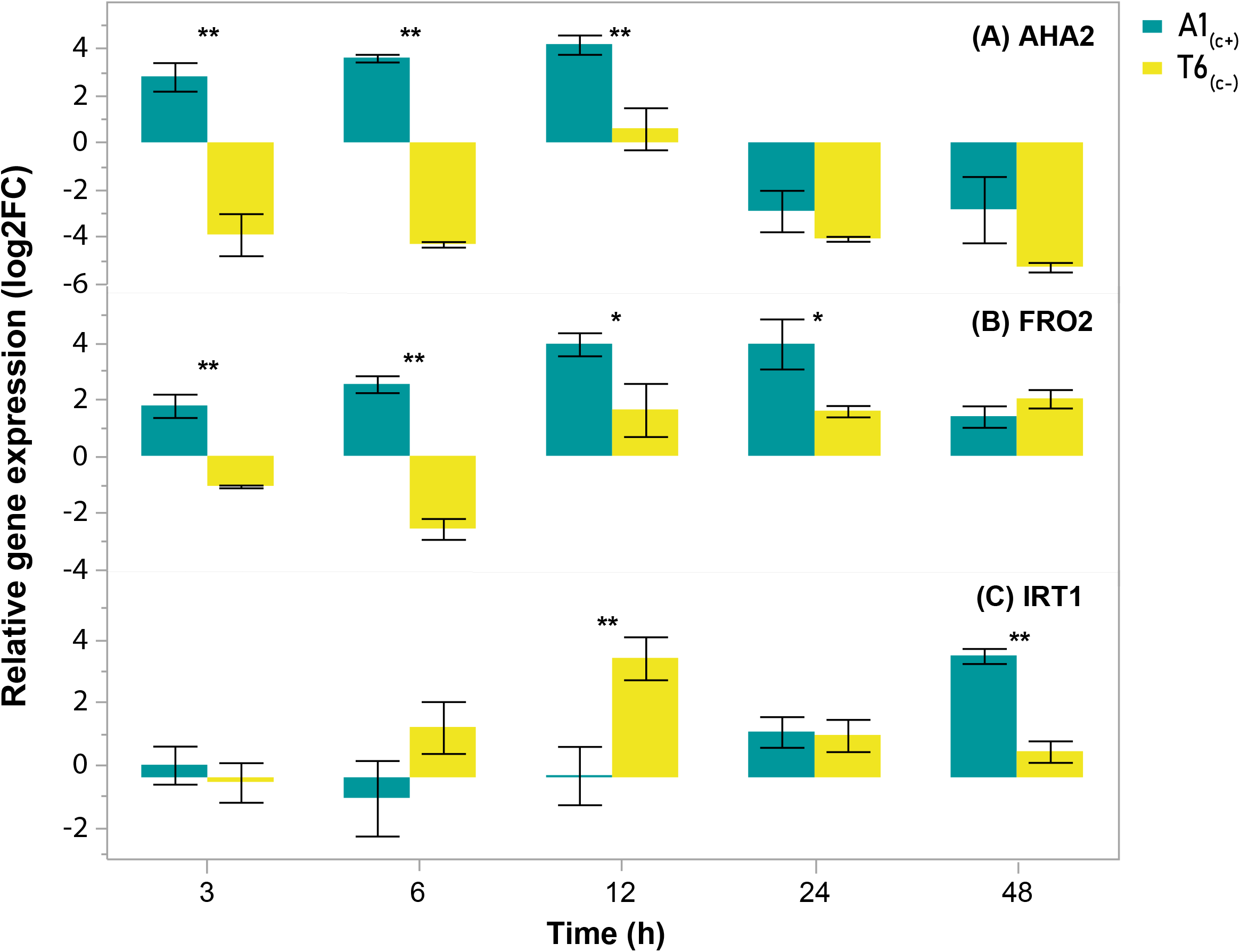
Root expression profiles of genes involved in iron acquisition in A1_(c+)_ and T6_(c−)_ under bicarbonate treatment. Relative expression (log2(FC)_Bic_ / log2(FC)_C_) of **(A)** AHA2, **(B)** FRO2, and **(C)** IRT1 (mean ± SE) in roots of A1_(C+)_ and T6_(C-)_ after 3 or 48h of bicarbonate exposure (N=6). Asterisks indicate significant differences (Student t-Test, adj. p-value < 0.05).

### 3.3 Relative expression of root S-type anion channel and aquaporin genes

In the view of the differential expression of *BCA4*.*1* in A1_(C+)_ and T6_(C-)_ and the role of this CA in the activation of *SLAC1* and stomatal closure (Wang et al., 2016), we compared the root expression profiles of *SLAH1* (SLAC1 homologue 1), *SLAH2* (SLAC1 homologue 2), and *SLAH3* (SALC1 homologue 3) in both demes under bicarbonate exposure (Figure 5). After 3 h exposure to bicarbonate both *SLAH1* and *SLAH3* were upregulated in A1_(C+)_ but not in T6_(C-)_. After 48 h SLAH1 was downregulated in A1_(C+)_ coincident with the downregulation of βCA4.1 in this deme. Only minor changes in SLAH2 were observed in both demes (Figure 5). According to TAIR, SLAH1 and SLAH3 have similarity to the SLAC1 protein involved in ion homeostasis in guard cells. Although, they are not expressed in guard cells, they can complement a slac1-2 mutant suggesting similar functions. So far, the S-type anion channel SLAH3 in roots of *A. thaliana* has been related to acidity sensing, regulation of nitrogen-potassium homeostasis and Cl-transport (Hedrich and Geiger, 2017; Lehmann et al., 2021; Liu et al., 2023). Bicarbonate-induced expression of GsSLAH3 has been observed in Glycine soja and overexpression of this gene in *A. thaliana* enhanced bicarbonate tolerance and the overexpressing lines accumulated higher nitrate concentrations (Duan et al., 2017). Our results give indirect support to the view that βCA4.1 may be involved in the bicarbonate tolerance of *A. thaliana* deme A1_(C+)_ by activating the transcription of S-type anion channels.

**Figure 5.**
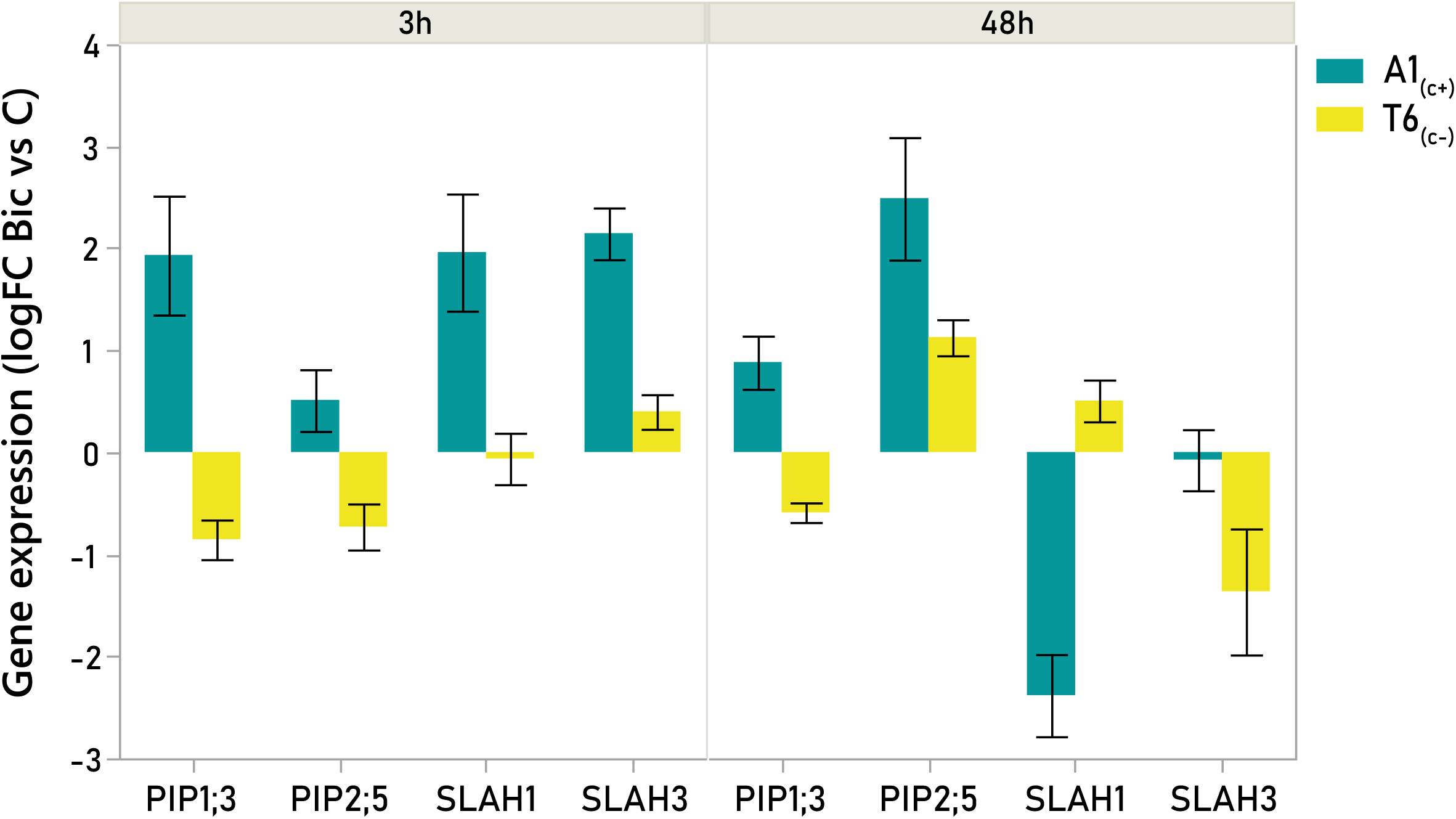
Root expression profiles of S-type anion channels and aquaporin genes in A1_(c+)_ and T6_(c−)_ under bicarbonate treatment based on cDNA microarray analysis. Relative expression (log2(FC)_Bic_ / log2(FC)_C_) of *PIP1;3, PIP2;5, SLAH1*, and *SLAH3* in roots of A1_(C+)_ and T6_(C)_ after 3 or 48h of bicarbonate exposure.

Moreover, in A1_(C+)_ bicarbonate caused a strong upregulation of *PIP1,3* and *PIP2,5* after 3- and 48-hours exposure, respectively (Figure 5). No activation of aquaporins was observed in T6_(C-)_. Besides water, aquaporins of the PIP1 type can transport CO_2_. Recently also *AtPIP2,5* has been shown to enhance more than 100-fold the CO_2_ entrance into yeast cells when co-expressed with carbonic anhydrase *AtCA1* (Israel et al., 2021). Our results support the view that a cooperation between carbonic anhydrase, aquaporin, and S-type anion channels may not only have a function in stomatal closure, but also in roots providing tolerance to high substrate bicarbonate.

### 3.4. βca4 mutant phenotyping

To get better insights into the possible role of βCA4 in bicarbonate tolerance, the performance of the βCA4;1 knockout mutant (βca4) exposed to bicarbonate was analysed in comparison to the WT (Col-0). No differences between mutant and wild type in rosette diameter and leaf chlorophyll concentrations were observed (data not shown). However, the mutant developed a lower number of leaves (Figure 6A) and higher root MDA content (Figure 6B). Mutant and WT also differed in root morphology (Figure 6C, D). Less lateral root formation and lower average root diameter were observed in the *βca4* mutant. The relatively small differences between the single loss-of-function mutant and WT phenotypes under bicarbonate exposure is probably due to the redundancy of the multiple CAs in *A. thaliana* (Bricks et al., 2006). Nonetheless, the influence of bicarbonate on root morphology in βca4, in comparison to WT, supports a specific role for βCA4.1 in iron acquisition. Enhanced root branching induced by Fe-deficiency may contribute to enhanced rhizosphere acidification and FRO2 activity favouring iron reduction (Jin et al., 2008). In fact, in the carbonate tolerant A1_(C+)_ the bicarbonate induced activation of BCA4.1 transcription (Figure 3) was accompanied by a higher enhancement of both *AHA2* and *FRO2* expression (Figure 4A, B) than in the carbonate sensitive T6_(C-)_ where βCA4.1 transcription was downregulated.

**Figure 6.**
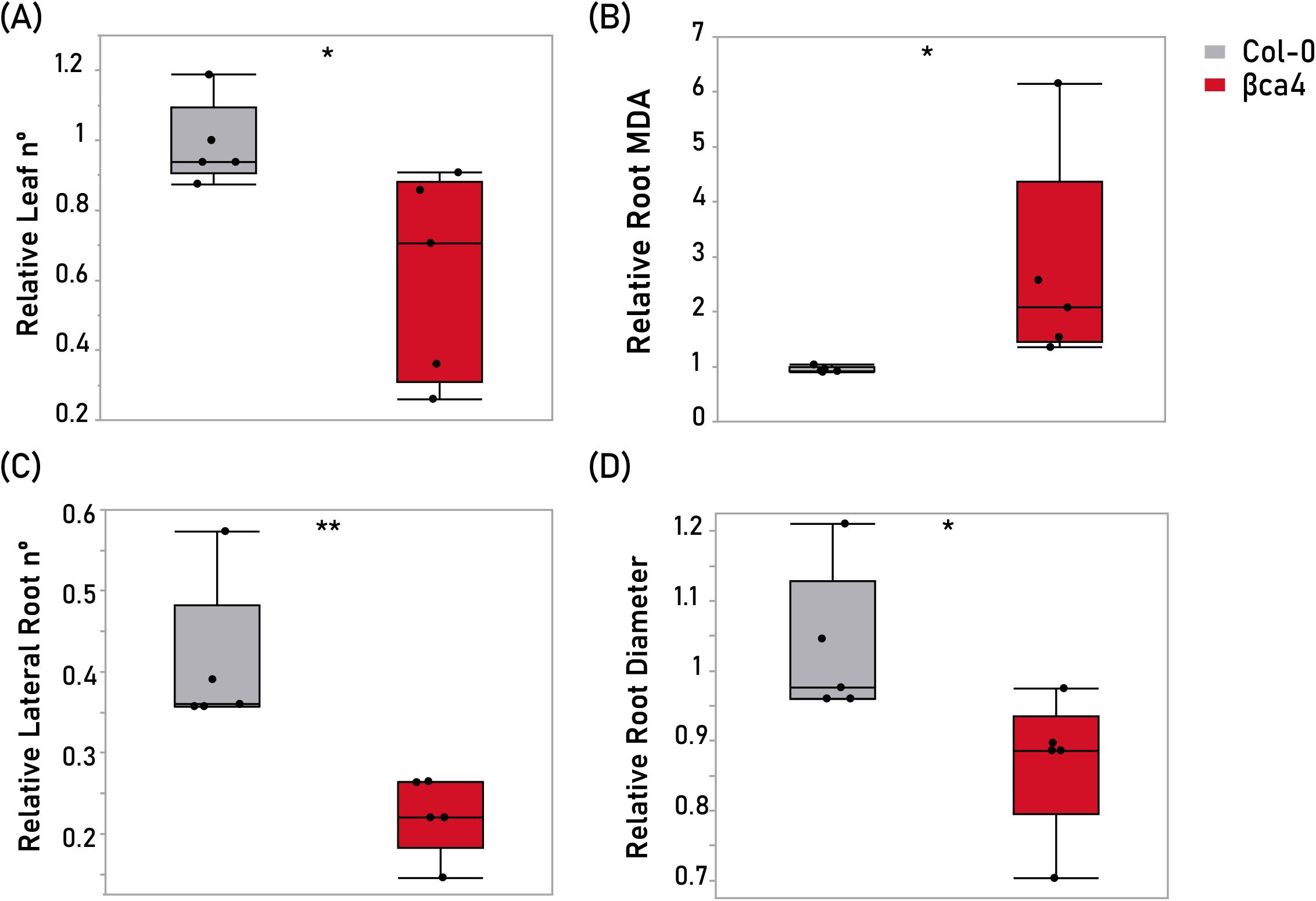
*BCA4* mutant phenotyping. Relative mean ± SE of **(A)** leaf number **(B)** root MDA, **(C)** lateral root number and **(D)** root diameter of the bca4;1 knockout mutant and Col-0 (WT) after 10 days of bicarbonate exposure (N=5). Asterisks indicate significant differences (Student t-Test, *: adj. p-value < 0.05; **: adj. p-value < 0.01).

## 4 Conclusions

The strong, bicarbonate-induced hyperpolarisation of the root cell plasma membrane may act as a quick (minutes) differential stress signalling mechanism in *A. thaliana* deme A1_(C+)_ with naturally evolved tolerance to bicarbonate. This finding adds a new, still poorly explored piece to the model of strategy 1 of Fe acquisition in plants exposed to alkaline soils.

The coincidence of higher expressions in A1_(C+)_ of β*CA4*.*1* with that of *AHA2* and *FRO2* along with the root phenotype of the *bca4* mutant provides circumstantial evidence for a role of βCA4.1 in the activation of these mechanisms involved in Fe acquisition under bicarbonate stress. In analogy to the already well described action of βCA4 in stomatal closure, where high internal HCO_3_ ^-^ concentrations activate aquaporins and anion channels, the early upregulation in A1_(C+)_ roots of βCA4.1, S-type anion channels SLAH3 and SLAH1, and PIP1,3 by bicarbonate suggests a cooperation of aquaporins and S-type anion channels in bicarbonate tolerance mediated by βCA4.1.

Further electrophysiological and molecular studies are required to see whether, and if so, how the initial membrane hyperpolarization is related to the subsequent (hours) upregulation of *βCA4*.*1, SLAH1, SLAH3, PIP1,2, AHA2*, and *FRO2*.

## AUTHOR CONTRIBUTIONS

CP, RT and SB conceived the study. LPM, MAC, and LEM performed cultivations, lab experiments and analysis. LPM and LR conducted electrophysiology experiments and data analysis. LPM, MAC, and SB performed the statistical and data analysis. CP and SB wrote the manuscript with input from all authors. All authors edited and approved the final manuscript.

## DATA AVAILABILITY STATEMENT

Data will be available on request.

## SUPPORTING INFORMATION

Supplementary Dataset S1. Primers list for qPCR analysis.

Supplementary Dataset S2. Statistical analysis of all the experiments conducted.

## REFERENCES

Bakker, E.G., Stahl, E.A., Toomajian, C., Nordborg, M., Kreitman, M. & Bergelson, J. (2006) Distribution of genetic variation within and among local populations of Arabidopsis thaliana over its species range. Molecular Ecology, 15(5), 1405–1418. 10.1111/j.1365-294X.2006.02884.x

Beck, J.B., Schmuts, H. & Schaal, B.A. (2008) Native range genetic variation in Arabidopsis thaliana is strongly geographically structures and reflects Pleistocene gacial dynamics. Moecular Ecology 17(3), 902–915. 10.1111/j.1365-294X.2007.03615.x

Bergelson, J., Stahl, E., Dudek, S. & Kreitman, M. (1998) Genetic variation within and among populations of Arabidopsis thaliana. Genetics, 148, 1311–1323. 10.1093%2Fgenetics%2F148.3.1311

Bricks, G.C., Osmont, K.S., Shindo, C., Sibout, R. & Hardtke, C.S. (2006) Unequal genetic redundancies in Arabidopsis – a neglected phenomenon? Trends in Plant Science 11(10), 492–498. 10.1016/j.tplants.2006.08.005

Demidchik, V., Bowen, H.C., Maathius, F.J.M., Shabala, S., Tester, M.A., White, P.J. & Davies, J. (2002) Arabidopsis thaliana root non-selective cation channels mediate calcium uptake and are involved in growth. The Plant Journal, 32, 799–808.

DiMario, R.J., Quebedeaux, J.C., Longstreth, D.J., Dassanayake, M., Hartman, M.M. & Moroney, J.V. (2016) The cytoplasmic carbonic anhydrase βCA2 and βCA4 are required for optimal plant growth at low CO2. Plant Physiology 171, 280–293. http://www.plantphysiol.org/cgi/doi/10.1104/pp.15.01990

DiMario, R.J., Clayton, H., Mukherjee, A., Ludwig, M. & Moroney J.V. (2017) Plant carbonic anhydrases: Structures, locations, evolution, and physiological roles. Molecular Plant 10(1), 30–46. doi: 10.1016/j.molp.2016.09.001.

DiMario, R.J., Machingura, M.C., Waldrop, G.L. & Moroney, J.V. (2018) The many types of carbonic anhydrases in photosynthetic organisms. Plant Science 268, 11–17. doi: 10.1016/j.plantsci.2017.12.002.

Duan, X., Yu, Y., Duanmu, H., Chen, C., Sun, X., Cao, L., & Zhu, Y. (2018) GsSLAH3, a Glycine soja slow type anion channel homolog, positively modulates plant bicarbonate stress tolerance. Physiologia Plantarum 164(2), 145–162. 10.1111/ppl.12683

Hassani, A., Azapagic, A. & Shokri, N (2021) Global predictions of primary soil salinization under changing climate in the 21st century. Nature Commununications, 12, 6663. 10.1038/s41467-021-26907-3

Hedrich, R. & Geiger, D. (2017) Biology of SLAC1-type anion channels-from nutrient uptake to stomatal closure. New Phytologist, 216(1), 46–61. 10.1016/j.biochi.2013.10.020

Hu, H, Rappel, W.J., Occhipinti, R., Ries, A., Böhmer, M., You, L., Xiao, C., Engineer, C.B., Boron, W.F. & Schroeder, J.I. (2015) Distinct cellular locations of carbonic anhydrases mediate carbon dioxide control of stomatal movements. Plant Physiology 169, 1168–1178. http://www.plantphysiol.org/cgi/doi/10.1104/pp.15.00646

Ignatova, L., Rudenko, N., Zhurikova, E., Borisova-Mubarakshina, M. & Ivanov, B. (2019) Metabolites, 9, 73. doi:10.3390/metabo9040073

Israel, D., Lee, S.H., Robson, T.M. & Zwlazek, J.J. (2022) Plasma membrane aquaporins of the PIP1 and PIP2 subfamilies facilitate hydrogen peroxide diffusion into plant roots. BMC Plant Biology, 22, 566. http://bmcplantbiol.biomedcentral.com/articles/10.1186/s12870-022-03962-6

Jin, C.W., Chen, W.W., Meng, Z.B. & Zheng S.J. (2008) Iron deficiency-induced increase of root branching contributes to the enhanced root ferruc chelate reductase activity. Journal of Integrative Plant Biology 50(12), 1557–1562. 10.1111/j.1744-7909.2008.00654.x

Kupriyanova, E.V., Pronina, N.A. & Los D.A. (2023) Adapting from low to high: An update to CO2-concentrating mechanisms of cyanobacteria and microalgae. Plants, 12, 1569. 10.3390/plants12071569

Lehmann, J., Jorgensen, M.E., Fratz, S., Müller, H.M., Kusch, J., Scherzer, S., & Maierhofer, T. (2021) Acidosis-induced activation of anion channel SLAH3 in the flooding-related stress response of Arabidopsis. Current Biology, 31(16) 3575–3585. 10.1016/j.cub.2021.06.018

Liu, B., Feng, C., Fang, X., Ma, Z., Xiao, C., Zhang, S.,. & He, K. (2023) The anion channel SLAH3 interacts with potassium channelsto regulate nitroge-potassium homeostasis and the membrane potential in Arabidopsis. The Plant Cell 35, 1259–1280. 10.1093/plcell/koad014

Livak, K.J. and Schmittgen, T.D., 2001. Analysis of relative gene expression data using real-time quantitative PCR and the 2-ΔΔCT method. Methods, 25(4), pp.402–408.

Martín-Barranco, A., Spielmann, J., Dubeaux, G., Vert, G. & Zelazny E. (2020). Dynamic control of the high affinity iron uptake complex in root epidermis cells. Plant Physiology 184, 1236–1250.

Momayyezi, M., McKown, D., Bell, S.C.S. & Guy, R.D. (2020) Emeging roles for carbonic anhydrase in mesophyll conductance and photosynhesis. The Plant Journal, 101, 831–844. doi: 10.1111/tpj.14638

Müller, W.E.G., Qiang, L., Schröder, H.C., Hönig, N., Yuan, D., Grebenjuk, V.A., Mussino, F., Giovine, M. & Wang X. (2014) Carbonic anhydrase: a key regulatory and detoxifying enzyme for Karst plants. Planta, 239, 213–229. DOI 10.1007/s00425-013-1981-2

Pérez-Martín L., Busoms S., Tolrà R., Poschenrieder C. (2021) Transcriptomics reveals fast changes in salicylate and jasmonate signaling pathways in shoots of carbonate tolerant Arabidopsis thaliana under bicarbonate exposure. International Journal of Molecular Sciences 22(3), 1226;

Poschenrieder, C., Fernández, J.A., Rubio, L., Pérez, L., Terés, J. & Barceló, J. (2018) Transport and use of bicarbonate in plants: current knowledge and challenges ahead. International Journal of Molecular Science, 19, 1352. 10.3390/ijms19051352

Raven, J.A., Giordano, M., Beardall, J., Maberly, S.C. Algal and aquatic plant carbon concentrating mechanisms in relation to environmental change. Photosynth Res. 2011 Sep;109(1-3, :281–96. doi: 10.1007/s11120-011-9632-6.

Riaz, N. & Guerinot M.L. (2021) All together now: regulation of the iron deficiency response. Journal of Experimental Botany 72(6), 2045–2055. doi:10.1093/jxb/erab003

Rubio, L., García, D., García-Sánchez, M. J., Niell, F. X., Felle, H. H., Fernández, J. A. (2017). Direct uptake of HCO3– in the marine angiosperm Posidonia oceanica (L.) Delile driven by a plasma membrane H+ economy. Plant Cell Environ. 40, 2820–2830. doi: 10.1111/pce.13057

Schmid, K.J., Törjék, O., Meyer, R., Schmuts, H., Hoffmann, M.H. & Altmann, T. (2006) Evidence for a large-scale population structure of Arabidopsis thaliana from genome-wide single nucleotide polymorhism markers. Theoretic and Applied Genetics, 112, 1104–1114. DOI 10.1007/s00122-006-0212-7

Scholl, S. T. May, and D. H. Ware, “Seed and Molecular Resources for Arabidopsis,” Plant Physiol., vol. 124, no. 4, pp. 1477–1480, 2000.

Stenøien, H.K., Fenster, C.B., Tonteri, A. & Savolainen, O. (2005) Genetic variability in natural populations of Arabidopsis thaliana from northern Europe. Molecular Ecology, 14, 137–148. 10.1111/j.1365-294X.2004.02359.x

Tagliavini, M. & Rombolà, A.D. (2001) Iron deficiency and chlorosis in orchard and vineyard ecosystems. European Journal of Agronomy, 15, 71–92. 10.1016/S1161-0301(01)00125-3

Terés J., Busoms S., Perez-Martin L., Luis-Villarroya A., Flis P., Álvarez-Fernández A., Tolrà R., Salt D. & Poschenrieder, C. (2019) Soil carbonate drives local adaptation in Arabidopsis thaliana. Plant Cell & Environment, 42, 2384–2398. DOI:10.1111/pce.13567

Tian, W., Hou, C., Ren, Z., Pan, Y., Jia, J., Zhang, H., Bai, F., Zhang, P., Zhu, H., He, Y., Luo, S., Li, L. & Luan S. (2015) A molecular pathway for CO2 response in Arabidopsis guard cells. Nature Communications, 6, 6057. DOI: 10.1038/ncomms7057

Vélez-Bermúdez, I.C. & Schmidt, W. (2023) plant strategies to mine iron from alkaline substrates. Plant and Soil 483, 1–25. 10.1007/s11104-022-05746-1

Wahba, M.M., Labib, F. & Zaghloul, A. (2019) management of calcareous soils in arid regions. International Journal of Pollution & Environmental Modelling, 2(5), 248–258.

Wang, M., Zhang, Q., Liu, F.C., Xie, W.F., Wang, G.D., Wang, J., Gao, Q.H. & Duan, K. (2014) Family-wide expression characterization of Arabidopsis beta-carbonic anhydrase genes using qRT-PCR and promoter: Gus fusions. Biochimie 97, 219–227. 10.1016/j.biochi.2013.10.020

Wang, N., Dong, X., Chen, Y., Ma, B., Yao, C., Ma, F., Liu, z. (2020) Direct and bicarbonate-induced iron deficiency differently affect iron translocation in kiwifruit roots. Plants 9(11), 1578. 10.3390/plants9111578

Weerasooriya, H.N., Longstreth, D.J., DiMario, R.J., Rosati, V., Cassel, B.A. 6 Moroney, J.V. (2024) Carbonic anhydrases in the cell wall and plasma membrane of Arabidopsis thaliana are required for optimal plant growth on low CO2. Frontiers in Plant Science, 11, 1267046. doi: 10.3389/fmolb.2024.1267046

Wu, H., Zhang, X., Giraldo J.P. & Shabala, S. (2018) It is not all about sodium: revealing tissue specificity and signalling roles of potassium in plant responses to salt stress. Plant and Soil, 431, 1–17. 10.1007/s11104-018-3770-y

Wu, Y.Y. & Xing, D.K. (2012) Effect of bicarbonate treatment on photosynthetic assimilation of inorganic carbon in two plant species of Moraceae. Photosynthetica 50(4), 587–594. DOI: 10.1007/s11099-012-0065-z

Wu, Y. & Wu, Y. (2022) The diversification of adaptive strategies for Karst-adaptable plants and the utilization of plant resources in Karst ecosystems. Agronomy, 13(8), 2135. 10.3390/agronomy13082135

Zhou, L., Xiang, X., Ji, D., Chen, Q., Ma, T., Wang, J. & Liu, C. (2024) A carbonic anhydrase, ZmCA4, contributes to photosynthetic efficiency and modulates CO2 signaling gene expression by interacting with aquaporin ZmPIP2,6 in maize. Plant and Cell Physiology 65(2), 243–258. 10.1093/pcp/pcad145

